# Impairment of hippocampal astrocyte-mediated striatal dopamine release and locomotion in Alzheimer’s disease

**DOI:** 10.1101/2023.10.29.564619

**Authors:** Benjamin B. Tournier, Kelly Ceyzériat, Aurélien M. Badina, Yessica Gloria, Aïda Fall, Quentin Amossé, Stergios Tsartsalis, Philippe Millet

**Author notes:** Corresponding author: Benjamin B. Tournier University Hospitals of Geneva Department of Psychiatry Avenue de la Roseraie, 64 1205 Geneva. **Abbreviations:** CNO: clozapine N-oxide; DREADD: Designer Receptors Exclusively Activated by Designer Drugs; Tg: TgF344-AD.

## Abstract

Clinical and translational research has identified deficits in the dopaminergic neurotransmission in the striatum in Alzheimer’s disease (AD) and this could be related to the pathophysiology of psychiatric symptoms appearing even at early stages of the pathology. We hypothesized that AD pathology in the hippocampus may influence dopaminergic neurotransmission even in the absence of AD-related lesion in the mesostriatal circuit. We thus chemogenetically manipulated the activity of hippocampal neurons and astrocytes in wild-type and hemizygous TgF344-AD (Tg) rats, an animal model of AD pathology. We assessed the brain-wide functional output of this manipulation using *in vivo* Single Photon Emission Computed Tomography to measure cerebral blood flow and D_2/3_ receptor binding. We also assessed the effects of the chemogenetic manipulations on astrocytic and microglial capacity to surround and phagocytize Aβ both locally and in the striatum. Our results show that acute and chronic neuronal and astrocytic stimulation induces widespread effects on the brain regional activation pattern, notably with an inhibition of striatal activation. In the TgF344-AD rats, both these effects were blunted. Chemogenetic stimulation in the hippocampus increased microglial density and its capacity to limit AD pathology, whereas these effects were absent in the striatum perhaps as a consequence of the altered connectivity between the hippocampus and the striatum. Our work suggests that hippocampal AD pathology may alter mesostriatal signalling and induce widespread alterations of brain activity. Neuronal and astrocytic activation may induce a protective, Aβ-limiting phenotype of microglia, which surrounds Aβ plaques and limits Αβ concentration more efficiently.

## Introduction

Alzheimer’s disease (AD) is a severe neurodegenerative condition characterized by progressive cognitive decline and functional disability. AD neuropathology involves the extracellular accumulation of insoluble forms of β amyloid peptide (Aβ) and intracellular accumulation of a hyperphosphorylated form of Tau protein forming neurofibrillary tangles (NFT). A third neuropathological hallmark in AD consists in “neuroinflammation” which describes phenotypic and functional alterations in glial cells, notably astrocytes and microglia^1–3^. *APOE*, the most important genetic risk factor for AD is predominantly expressed in astrocytes, which are activated early in the disease process and may phagocytize Aβ plaques. Astrocyte function and dysfunction are intimately related to microglial states in AD^2,4–10^ and some studies suggest that astrocytes may show signs of activation even before microglia^11,12^. Microglial cells also have a major role in the disease pathophysiology phagocytizing Aβ thus limit the lesion burden in the brain. In addition, multiple gene associated with the genetic risk for AD are predominantly expressed in microglia, whereas. Overall, glial cells are important targets for the understanding of AD and its manifestations and for the development of therapeutic strategies.

AD is also associated to behavioural and psychological symptoms of dementia, a clinical entity that includes multiple syndromes, some of which, such as psychosis, have been associated to dopaminergic dysfunction^13–15^. In addition, these symptoms may also appear early in the disease process, even before the emergence of overt cognitive symptoms. Dopaminergic circuitry dysfunction has been shown in AD though clinical neuroimaging studies^16–19^, whereas AD animal model studies also indicate a dopaminergic dysfunction, even at early stages of the pathology^20–22^. The occurrence of dopaminergic dysfunction at stages of the disease where widespread neuropathological lesions are not yet present raises the hypothesis that areas such as the hippocampus, which is affected early in AD, may influence the function of the mesostriatal circuit and thus induce a dopaminergic dysfunction.

Previous work has indicated that optogenetic manipulation of hippocampal neuron activity can induce dopamine release in the striatum^23^. One may thus hypothesize that early AD pathology in the hippocampus may indirectly influence striatal dopaminergic activity via a circuit from the hippocampus to the ventral tegmental area (VTA) where the dopaminergic connections to the striatum stem from.

For this reason, we employed a chemogenetic strategy using Designer Receptors Exclusively Activated by Designer Drugs (DREADD) to manipulate the activity of hippocampal neurons and astrocytes in wild-type and hemizygous TgF344-AD (Tg) rats, an animal model of AD pathology. We assessed the brain-wide functional output of this manipulation using (i) *in vivo* [^99m^Tc]HMPAO Single Photon Emission Computed Tomography (SPECT), an approach measuring cerebral blood flow as a proxy for regional brain activity and (ii) [^123^I]IBZM SPECT, an approach assessing D_2/3_ receptor binding, which may measure both the D_2/3_ receptor concentration and its occupancy by dopamine (a proxy for dopamine release)^24,25^. We also assessed the effects of the hippocampal neuron and astrocyte manipulation on astrocyte and microglial activity and capacity to surround and phagocytize Aβ both locally and in the striatum. Our results point to an impact of hippocampal AD pathology on mesostriatal dopaminergic signalling, whereas local stimulation of neurons and astrocytes may enhance the capacity of microglia to limit AD pathology.

## Materials and Methods

### Experimental design and CNO treatment

At the age of 5 months, the animals were bilaterally injected in the hippocampus with one of the DREADD or AAV. After one month of recovery, the animals received two injections of clozapine N-oxide (CNO, 3mg/kg i.p.) to measure its acute effect on dopamine and regional brain blood flow on day 1 using SPECT imaging, and locomotor activity on day 2. The CNO was then given by drinking water (0.015 mg/ml) for 28 days. D_2_R density and regional brain blood flow were measured again by SPECT imaging on day 27 and locomotor activity on day 28, at the age of 7 months. Euthanasia were carried out on day 28 for *postmortem* investigations. All animals received the CNO treatment.

### Animals

Male WT and hemizygote Fisher 344 (TgF344-AD, APPswe and PS1ΔE9 transgenes) rats aged 5 months at the start of the experiment were housed in a 12h light-dark cycle with food and water *ad libitum*. All experimental procedures were approved by the Ethics Committee for Animal Experimentation of the Canton of Geneva, Switzerland. Data are reported in accordance with Animal Research: Reporting In Vivo Experiments (ARRIVE) guidelines.

### Injection of DREADD and AAV

All viruses were produced by the Viral Vector Facility (Zurich, Switzerland) at a minimum concentration of 1.7 x 10^12^ vg/ml. For chemogenetic manipulations, rats were bilaterally injected with 2 µl of ssAAV-8/2-mCaMKIIa-hM3D(Gq)_mCherry-WPRE-hGHp(A) or ssAAV-9/2-hGFAP-hM3D(Gq)_mCherry-WPRE-hGHp(A) to activate neurons and astrocytes, respectively. Controls animals received ssAAV-8/2-mCaMKIIα-mCherry-WPRE-hGHp(A) or ssAAV-9/2-hGFAP-mCherry-WPRE-hGHp(A). Under isoflurane anesthesia and buprenorphine (Temgesic), injections were made into the hippocampus (AP: -4.8mm, Lat:+/-3mmm, V:-3.2 mm). After injecting at a rate of 0.2 µl/min, the syringe was left in place for 2 min before withdrawal.

### SPECT acquisition

One month after stereotactic injection, 40-min SPECT image acquisition was initiated 1h50, 1h20 and 15 min after injection of (CNO, 3mg/kg i.p.), [^125^I]IBZM (26.87 ± 5.7 MBq, i.v.) and [^99m^Tc]HMPAO (66.47 ± 15.7 MBq, i.v.), in agreement with previous studies^24^. [^125^I]IBZM was synthesized as previously described^24^ and [^99m^Tc]HMPAO according to the manufacturer protocol (Ceretec, GE Healthcare). Images were reconstructed using a POSEM (0.4mm voxels, four iterations, six subsets) approach, with radioactive decay correction applied. Image processing was done using PMOD software (PMOD3.5, PMOD Technologies Ltd, Zurich, Switzerland) and were analyzed firstly using the VOI template integrated to PMOD^26^ to extract the radioactivity from each brain VOI, and the cerebellum and the whole brain, which were used as reference regions for the specific binding ratio (SBR) calculation for [^125^I]IBZM and [^99m^Tc]HMPAO, respectively. Differences between controls and DREADD activated groups were estimated as (100*SBR_DREADD_)/(SBR_CONTROL_)−100. In addition, statistical analyses were performed at the voxel level using the SPM12 software (Wellcome Trust Centre for Neuroimaging, UCL, London, UK) and the Small Animal Molecular Imaging Toolbox^25^ (SAMIT1.3, SAMIT, Groningen, Netherlands) in Matlab (R2019, Mathworks Inc, USA).

### Locomotor activity

The apparatus consists of 4 square boxes of 45x45x40 cm overlooked by a digital camera. The distance travelled was automatically analysed by the Noldus software (EthoVision, Noldus). The day following the first scan acquisition, animals received a second i.p. injection of CNO and their 30-min locomotor response was recorded 30 min after the injection. The day following the second scan acquisition (i.e. after 43 weeks of chronic CNO treatment, 0.015mg/ml in drinking water), the locomotor activity was recorded for 30 min.

### Euthanasia

Under isoflurane anesthesia, an intracardiac saline perfusion was performed before brain removal. One hemisphere was immersed in formalin for 24 hours and then in a sucrose solution of increasing concentration before being frozen and cut with a cryostat (30 μm sections) for carrying out immunofluorescence. The other hemisphere was dissected to isolate the hippocampus and the striatum for protein extraction. The samples were frozen in liquid nitrogen and then stored at -80°C.

### Immunofluorescence

Serial brain sections (35 μm) covering the striatum or the hippocampus were realized with a cryostat. Sections were incubated with primary antibodies overnight at 4°C in 1% BSA/PBS 0.1M/0.3% Triton X-100. The antibody list is as follows: anti-IBA1 (1/600, Rabbit; Wako), anti-GFAP-Cy3 (1/1000, Sigma), anti-GFAP (1/500, mouse, Thermofisher), anti-CHERRY (1/500, rabbit, Thermofisher), anti-Neun (1/500, guinea pig; Synaptic System), and anti-FOS (1/500, mouse, Abcam). Three 5-min PBS 0.1M washes was performed before incubation for 1h at room temperature in 1% BSA/PBS 0.1M/0.3% Triton X-100 with the appropriate secondary Alexa-fluor conjugated antibodies (1/200, Invitrogen). Sections were then washed (3x 5-min PBS 0.1M), immerged in Sudan black (0.1% in 70% EtOH, 10 min), washed (3x 5-min PBS 0.1M), and counterstained with DAPI (10 min) before being washed (3x 5-min PBS 0.1M) and mounted on gelatin slides and coated with FluorSave (Calbiochem). Images of the whole section were taken using Axioscan.Z1 (Zeiss) at 10x and analyzed using ImageJ software. Confocal images (40x, 2 µm steps) were used for the measure of MXO4-IBA1 and MXO4-GFAP colocalization in the hippocampus. % of MXO4 covered by IBA1 and GFAP was measured using ImageJ.

### Protein extraction and ELISA

Hippocampus and striatum samples were sonicated after immersion in a solution of triton (50mM Tris HCl, 150mM NaCl, 1%Triton x100, protease and phosphatase inhibitors 1x, pH=7.4). After centrifugation (20 000 g, 20 min, 4°C), the supernatants collected form the triton soluble fraction (Tx) and were stored at -80°C. The pellet was taken up with a solution of guanidine (5M guanidine, 50mM Tris HCl, protease, and phosphatase inhibitors 1x, pH=8). After gentle agitation (3h, 4°C) and centrifugation (20 000 g, 20 min, 4°C), the supernatants collected form the guanidine soluble fraction (Gu). Amyloid Aβ40 and Aβ42 in the hippocampus and striatum and D_2_R in the striatum were quantified by ELISA with the following kits: Aβ40 human ELISA kit (Thermofisher); Aβ42 human ultrasensitive ELISA Kit (Thermofisher), and D_2_R (Mybiosource). The procedure was performed according to the instructions of the kits. Briefly, the samples were exposed for 2-3 hours to the specific antibody attached to the wells of a 96-well plate. After washing, the primary and then the secondary antibody were introduced into the wells for 30 min before adding the stabilized chromogen (30 min) and the stop solution. The OD reading was performed at 450 nm in the presence of a standard curve, specific to each kit.

### Statistics

A sample size analysis with the graphical Douglas Altman’s nomogram^27^ was performed considering our previous observations on SPECT-D2R measurement in the Tg rats^21^. Animals were randomly allocated to the different treatment groups. Comparisons were made by two-way ANOVA with Dunnet’s or Fisher’s LSD *posthoc* tests, depending on the type of comparisons desired.

## Results

### Selective expression of DREADD in neurons or astrocytes

To directly modulate the activity of neurons or astrocytes, we used designer receptors exclusively activated by designer drugs (DREADDs) delivered by adeno-associated virus (AAV) vectors. Bilateral injections into the hippocampus of CNO-activable DREADD viruses was performed. AAV contained either a *CamKII* or a *Gfap* promoter-driven gene to drive the expression in neurons and astrocytes, respectively. The specificity of the DREADD expression system was validated using immunofluorescence. As shown in the representative example in Fig. 1, the expression of the mCherry reporter gene was colocalized with NEUN but not with GFAP, confirming the neuronal specificity of the AAV-*CamKII*. In addition, the AAV-*Gfap* was selectively expressed in astrocytes and not in neurons.

**Figure 1.**
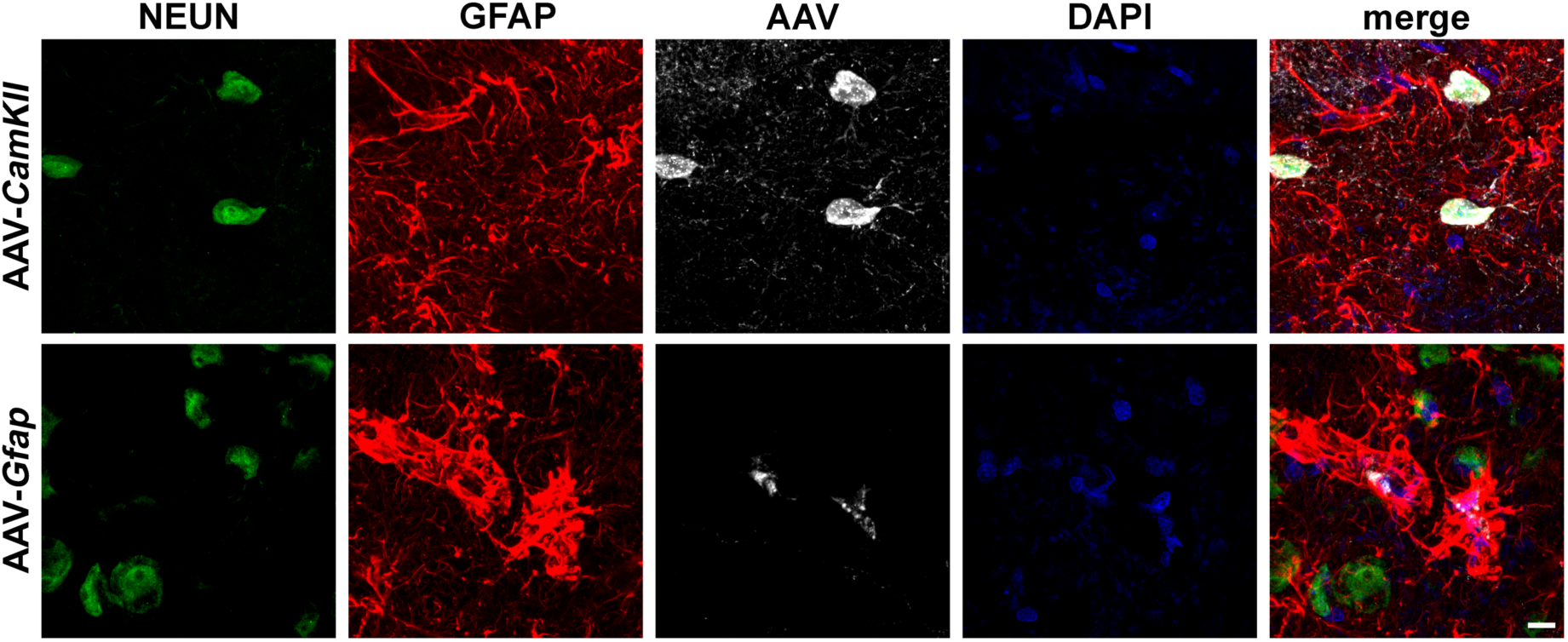
Cell-type specificity of AAV-*CamKII* and AAV-*Gfap* expression. Representative example of the cell identity of the AAV-*CamKII* and AAV-*Gfap*. Scale bar: 10µm

### Acute DREADD activation decreases IBZM binding in WT and, to a lesser extent, in Tg

Optogenetic activation of dorsal hippocampal neurons has been shown to induce dopamine release in the striatum^28^. An indirect measurement of dopamine release can be obtained by *in vivo* imaging using the measurement of D_2/3_R radioligand binding inhibition^29,30^ including with [^125^I]IBZM^21^. However, the impact of hippocampal astrocyte stimulation as well as the sensitivity of Tg animals remains unknown. We therefore performed SPECT acquisition with [^125^I]IBZM in response to i.p. injection of CNO (3mg/kg) in WT and Tg animals previously injected bilaterally into the hippocampus with neuron- or astrocyte-specific DREADD, or control AAV viruses (that neither induce activation of infected cells nor any response to CNO). A representative example of scan is shown in Fig. 2A. CPu showed greater binding of [^125^I]IBZM than Acc, in control WT and control Tg, with no differences between strains, which is consistent with the striatal D_2/3_R mapping (Fig. 2B, two-way ANOVA, main effect of genotype: F1,24=1.38, p>0.05; main effect of area: F1,24=47.48, p<0.0001; interaction: F1,24=0.11, p>0.05).

**Figure 2.**
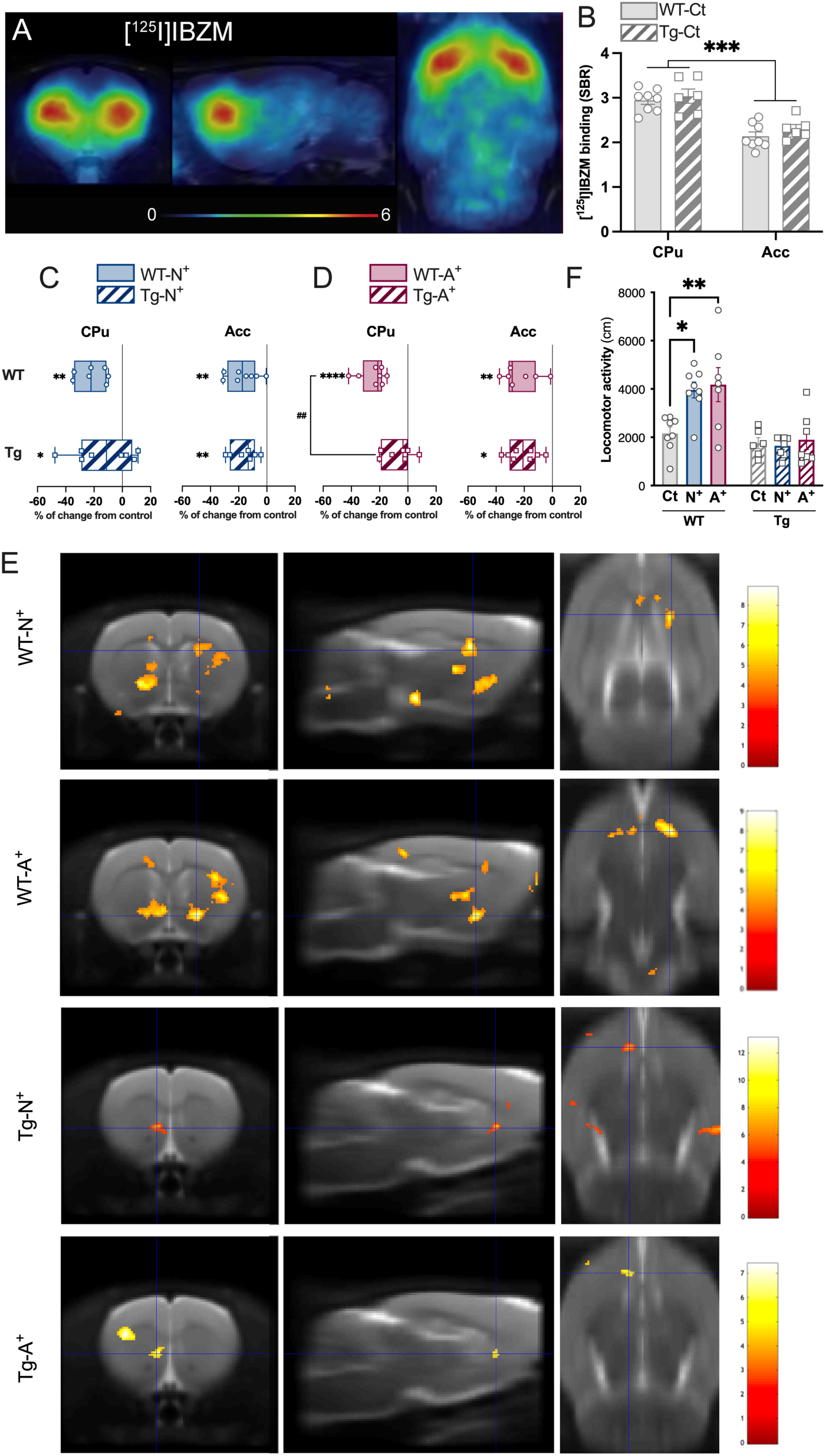
Acute CNO stimulation of hippocampal astrocytes induces a higher displacement of [^125^I]IBZM binding in CPu in WT than in TgF344-AD rats. **A.** Mean parametric maps of [^125^I]IBZM SBR in WT animals injected with control AAV after CNO injection at the level of the striatum. Images were coregistered to the MRI atlas in the coronal (left), sagittal (center), and horizontal (right) planes. Color bar indicates the specific binding ratio (SBR) from 0 to 3.5. **B.** Quantitative analysis of [^125^I]IBZM SBR in CPu and Acc in WT and Tg control animals (WT-Ct and Tg-Ct, respectively). **C.** Representation of the % of change of [^125^I]IBZM SBR in CPu and Acc of WT and Tg animals injected with AAV-N^+^ (WT-N^+^ and Tg-N^+^) or AAV-A^+^ (WT-A^+^ and Tg-A^+^) compared to their respective WT-Ct or Tg-Ct groups, in response to acute CNO. Two-way ANOVA analysis with the Fisher’s LSD posthoc test. *p<0.05, **p<0.01, ****p<0.0001 as compared to the respective Ct group and ##p<0.01 as compared to the WT-A^+^ group. **E.** Voxel-voxel group comparison using SAMIT (p<0.001). **F.** Quantification of the locomotor activity in response to acute CNO.

CNO-induced neuron-specific DREADD activation decreased [^125^I]IBZM binding in the caudate/putamen in a similar manner in WT and Tg rats (Fig. 2C, two-way ANOVA, main effect of AAV: F1,25=11.88, p=0.002; main effect of genotype: F1,25=1.05, p>0.05; interaction: F1,25=0.19, p>0.05). Neuronal activation also induced a decrease in [^125^I]IBZM binding in the accumbens similar in WT and Tg rats (Fig. 2C, two-way ANOVA, main effect of AAV: F1,25=24.49, p<0.0001; main effect of genotype: F1,25=3.04, p>0.05; interaction: F1,25=0.0003, p>0.05).

CNO-induced astrocyte-specific DREADD activation decreased [^125^I]IBZM binding in the caudate/putamen in WT but not in Tg rats (Fig. 2D, two-way ANOVA, main effect of AAV: F1,25=19.63, p<0.002; main effect of genotype: F1,25=7.07, p=0.013; interaction: F1,25=3.36, p=0.078). Astrocytic activation also induced a decrease in [^125^I]IBZM binding in the accumbens that was similar between WT and Tg rats (Fig. 2D, two-way ANOVA, main effect of AAV: F1,25=16.21, p=0.0005; main effect of genotype: F1,25=3.39, p=0.077; interaction: F1,25=0.0005, p>0.05).

Statistical comparison of the effect of astrocytes/neurons activation on [^125^I]IBZM binding was performed at the voxel level using SPM. Astrocyte and neuronal activation decreased [^125^I]IBZM binding in striatal voxels in WT animals and, to a lesser extent, in Tg animals (Fig. 2E).

Dopamine release in the striatum is known to stimulate locomotor activity. Thus, we next investigated the consequences of the DREADD-induced DA release on locomotion. Stimulation of hippocampal neurons and astrocytes induced a hyper-locomotion in WT animals that is absent in Tg (Fig. 2F, two-way ANOVA, main effect of AAV: F2,38=4.25, p=0.021; main effect of genotype: F1,38=28.48, p<0.0001; interaction: F2,38=3.98, p=0.027).

### Acute DREADD activation alters HMPAO accumulation in WT but not in Tg

The activity of brain areas is associated with cerebral blood flow which can be measured by *in vivo* imaging with [^99m^Tc]HMPAO. As the hippocampus is connected to many brain structures, a change in its activity is likely to lead to alterations in the activity of other brain regions. To test this idea and to measure the presence of a connectivity deficit in Tg vs WT animals, we performed a SPECT acquisition with [^99m^Tc]HMPAO (simultaneously with [^125^I]IBZM). The analysis covers the dorsal and ventral striatum, the hippocampus as well as all other brain regions, as shown in the representative example Fig. 3A. [^99m^Tc]HMPAO accumulation is dependent on brain regions but did not differ between WT control and Tg control rats (Supplemental Table 1, two-way ANOVA, main effect of genotype: F1,360=1.3, p>0.05; main effect of area: F29,360=76.5, p<0.0001; interaction: F29,360=1.56, p=0.035. The Šidák multiple comparisons test did not reveal any genotype difference regardless the brain area).

**Figure 3.**
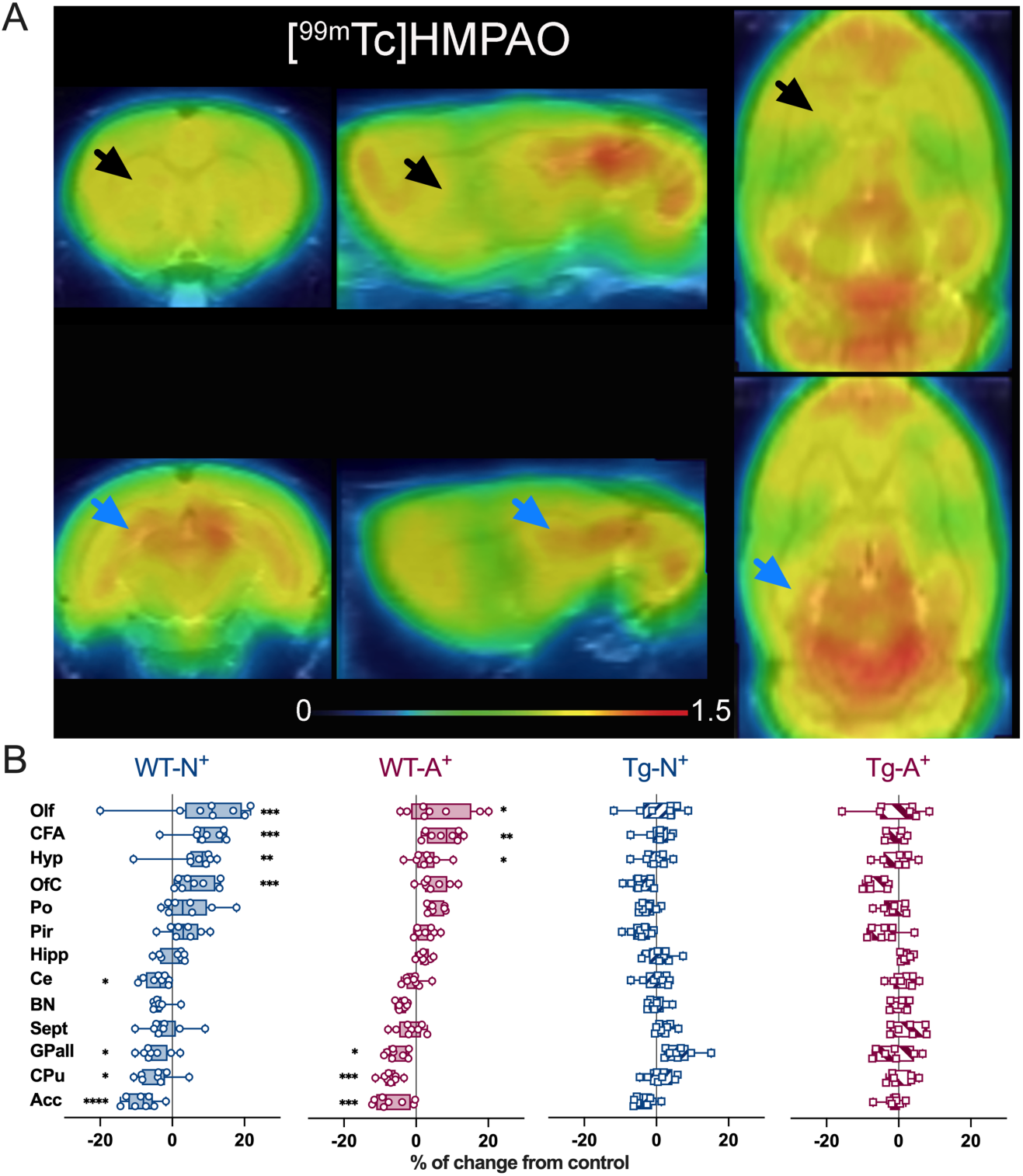
Acute CNO stimulation of hippocampal astrocytes and neurons induces a reduction in [^99^Tc]HMPAO binding in striatum in WT but not in TgF344-AD rats. **A.** Mean parametric maps of [^99^Tc]HMPAO whole-brain ratio in WT animals injected with control AAV after CNO injection at the level of the striatum (black arrows) and hippocampus (blue arrows). Images were coregistered to the MRI atlas in the coronal (left), sagittal (center), and horizontal (right) planes. Color bar indicates the whole-brain ratio from 0 to 1.5. **B.** % of change of [^99^Tc]HMPAO whole-brain ratio in response to acute CNO in selected brain areas of WT and Tg animals injected with N^+^ or A^+^ as compared to Ct groups. Two-way ANOVA analysis with the Dunnett’s multiple comparisons posthoc test. *p<0.05, **p<0.01, ***p<0.001, ****p<0.0001 as compared to the respective Ct group.

Acute stimulation of hippocampal neurons or astrocytes induces changes in cerebral blood flow in WT animals (Fig. 3B, two-way ANOVA, main effect of AAV: F2,630=1.67, p>0.05; main effect of area: F29,630=170.8 p<0.0001; interaction: F58,630=2.42, p<0.0001). Dunnett’s *posthoc* analysis with correction for multiple comparisons indicates that stimulation of neurons induced an increase in cerebral blood flow in the olfactory nucleus, the associative area of the frontal cortex, the hypothalamus and the orbitofrontal cortex. Conversely, a significant decrease is measured in the main dopaminergic regions (accumbens, caudate/putamen, and globus pallidus) and the cerebellum. Astrocyte stimulation induced an increase in cerebral blood flow in the olfactory nucleus, the associative area of the frontal cortex and the hypothalamus, and a significant decrease in the accumbens, the caudate/putamen and the globus pallidus. In contrast, acute stimulation of hippocampal neurons or astrocytes did not induce any change in cerebral blood flow in Tg (Fig. 3B, two-way ANOVA, main effect of AAV: F2,540=0.42, p>0.05; main effect of area: F29,540=115.6 p<0.0001; interaction: F58,540=0.88, p>0.05).

### Chronic DREADD activation decreases dopamine D2 receptors and alters HMPAO binding in WT but not in Tg

Many pathophysiological systems show different responses to acute and chronic stimulation. To determine the effects of chronic stimulation of hippocampal neurons or astrocytes on brain activity, CNO administration (15.87 mg/l) was conducted for 4 weeks (including the day of SPECT examination). Chronic stimulation of hippocampal neurons did not induce changes in [^125^I]IBZM binding in the caudate/putamen in WT and Tg rats (Fig. 4A, two-way ANOVA, main effect of AAV: F1,25=1.69, p>0.05; main effect of genotype: F1,25=6.82, p=0.015; interaction: F1,25=1.82, p>0.05). In contrast, chronic stimulation of neurons induced a decrease in [^125^I]IBZM binding in the accumbens in WT rats but not in Tg rats (Fig. 4B, two-way ANOVA, main effect of AAV: F1,25=1.1, p>0.05; main effect of genotype: F1,25=7.38, p=0.018; interaction: F1,25=7.44, p=0.011).

**Figure 4.**
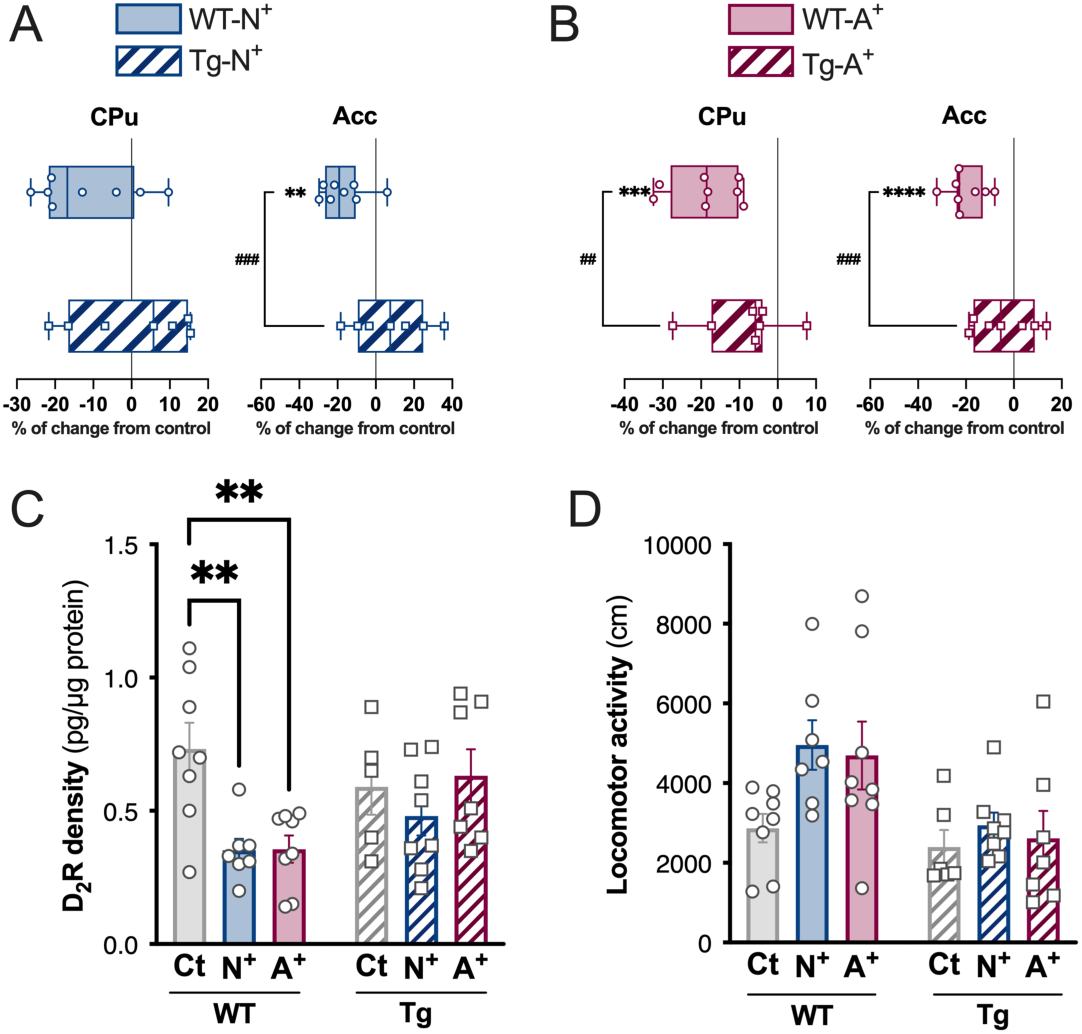
Chronic CNO stimulation of hippocampal astrocytes induces a reduction of D_2_R in striatum in WT but not in TgF344-AD rats. **A-B.** % of change of [^125^I]IBZM SBR in CPu and Acc of WT and Tg animals injected with AAV-N^+^ (WT-N^+^ and Tg-N^+^) or AAV-A^+^ (WT-A^+^ and Tg-A^+^) compared to their respective WT-Ct or Tg-Ct groups, in response to chronic CNO. 2-way ANOVA analysis with the Fisher’s LSD posthoc test. **p<0.01, ***p<0.001,****p<0.0001 as compared to the respective Ct group and #p<0.05, ##p<0.01 as compared to the respective WT-treated group. **C.** Quantitative analysis of D_2_R density in the striatum. Two-way ANOVA analysis with the Fisher’s LSD posthoc test. **p<0.01 as compared to the respective Ct group. **D.** Locomotor response to chronic CNO.

The chronic stimulation of hippocampal astrocytes induced a decrease in [^125^I]IBZM binding in the caudate/putamen and accumbens in WT rats but not in Tg rats (two-ay ANOVA caudate/putamen, main effect of AAV: F1,25=15.25, p=0.0006; main effect of genotype: F1,25=8.97, p=0.006; interaction: F1,25=1.94, p=0.17; two-way ANOVA accumbens, main effect of AAV: F1,25=14.84, p=0.0007; main effect of genotype: F1,25=6.85, p=0.014; interaction: F1,25=6.94, p=0.014).

A dowregulation of D_2/3_R has been repeatedly observed in response to chronic D_2/3_R stimulation. Thus, decreased *in vivo* binding of [^125^I]IBZM in response to chronic stimulation of hippocampal neurons and astrocytes may reflect dopamine release and/or decreased D_2/3_R concentration. To determine whether a reduction in D_2/3_R density was present, we performed a *postmortem* assay of striatal D_2_R quantification by ELISA. Chronic stimulation of hippocampal neurons and astrocytes induces a reduction in D_2_R density in WT animals that is absent in Tg, suggesting that the reduction in binding of [^125^I]IBZM is at least partly related to the decrease in D_2_R density (Fig. 4C, two-way ANOVA, main effect of AAV: F2,37=4.55, p=0.017; main effect of genotype: F1,37=1.73, p=0.19; interaction: F2,37=3.26, p=0.049). To measure the impact of D_2_R changes on DA-mediated behavior, we measured locomotion. The increase in locomotion that was observed in WT in response to acute stimulation of hippocampal neurons and astrocytes did not persist following chronic stimulation, when D_2_R decrease occurred. The locomotor activity of Tg rats was insensitive to chronic neurons/astrocytes stimulation (Fig. 4D, two-way ANOVA, main effect of AAV: F2,38=2.69, p=0.08; main effect of genotype: F1,38=10.05, p<0.003; interaction: F2,38=1.16, p>0.05).

Chronic stimulation of hippocampal neurons or astrocytes induces a change in cerebral blood flow in WT animals (Fig. 5, two-way ANOVA, main effect of AAV: F2,630=3.71, p=0.025; main effect of area: F29,630=109.7 p<0.0001; interaction: F58,630=2.92, p<0.0001). *Posthoc* analysis shows that neuronal stimulation induced an increase in cerebral blood flow in the olfactory nucleus, the associative area of the frontal cortex, the hypothalamus and the orbitofrontal cortex. Conversely, a significant decrease was measured in the accumbens, the caudate/putamen and the globus pallidus. Astrocyte stimulation induces an increase in cerebral blood flow in the olfactory nucleus, the associative area of the frontal cortex, the hypothalamus, the orbitofrontal cortex, the piriform cortex and the pons, and a significant decrease in the accumbens, the caudate/putamen, the bed nucleus of the stria terminalis and the cerebellum.

**Figure 5.**
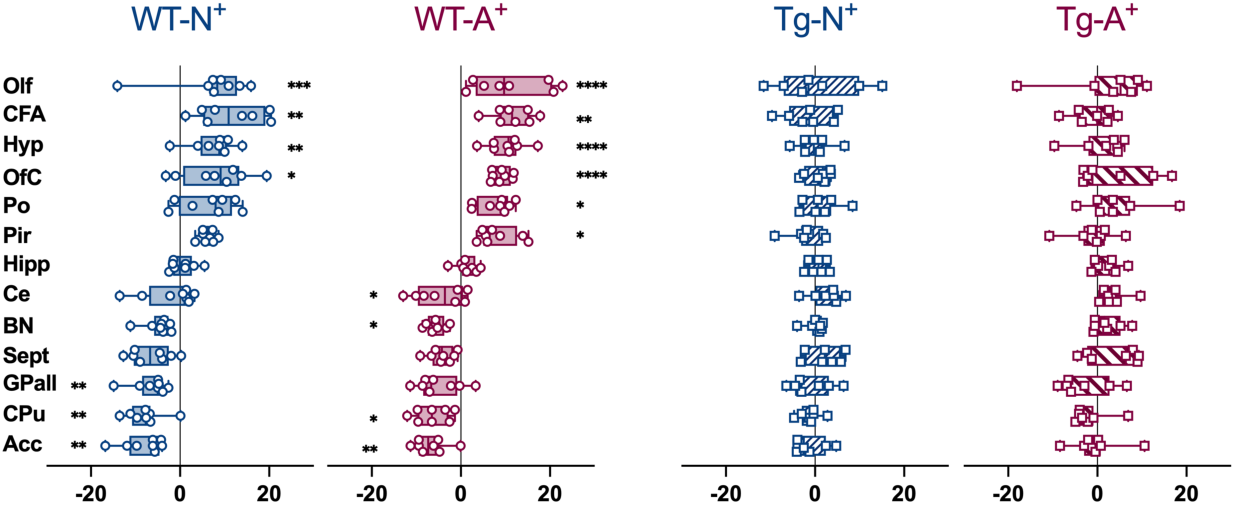
Chronic CNO stimulation of hippocampal astrocytes and neurons induces a reduction in [^99^Tc]HMPAO binding in striatum in WT but not in TgF344-AD rats. % of change of [^99^Tc]HMPAO whole-brain ratio in response to chronic CNO in selected brain areas of WT and Tg animals injected with N^+^ or A^+^ as compared to Ct groups. Two-way ANOVA analysis with the Dunnett’s multiple comparisons posthoc test. *p<0.05, **p<0.01, ***p<0.001, ****p<0.0001 as compared to the respective Ct group.

In contrast, chronic stimulation of hippocampal neurons or astrocytes did not induce any change in cerebral blood flow in Tg (Fig. 5, two-way ANOVA, main effect of AAV: F2,510=0.45, p>0.05; main effect of area: F29,510=102.3 p<0.0001; interaction: F58,510=0.9, p>0.05).

At voxel level, chronic astrocyte and neuronal activation reduced the [^99m^Tc]HMPAO binding in voxels from the striatum in WT animals. No changes were observed in Tg.

### Chronic DREADD activation decreases guanidine-soluble amyloid

To determine the ameliorative effects of chronic stimulation of hippocampal neurons or astrocytes on amyloid accumulation, we quantified triton- (Tx-) and guanidine- (Gu-) soluble forms to reveal low or high aggregated forms of Aβ40 and Aβ42. In the hippocampus, the chronic stimulation of neurons or astrocytes has no effect on Tx-Aβ40 and Tx-Aβ42 (Fig. 6A, one-way ANOVA Tx-Aβ40, main effect of treatment: F2,13=0.37, p>0.05; Tx-Aβ42, main effect of treatment: F2,18=0.23, p>0.05), but, in contrast, induced a decrease in Gu-Aβ40 and Gu-Aβ42 forms (Fig. 6B, one-way ANOVA Gu-Aβ40, main effect of treatment: F2,12=5.47, p=0.020; Gu-Aβ42, main effect of treatment: F2,13=5.34, p=0.02).

**Figure 6.**
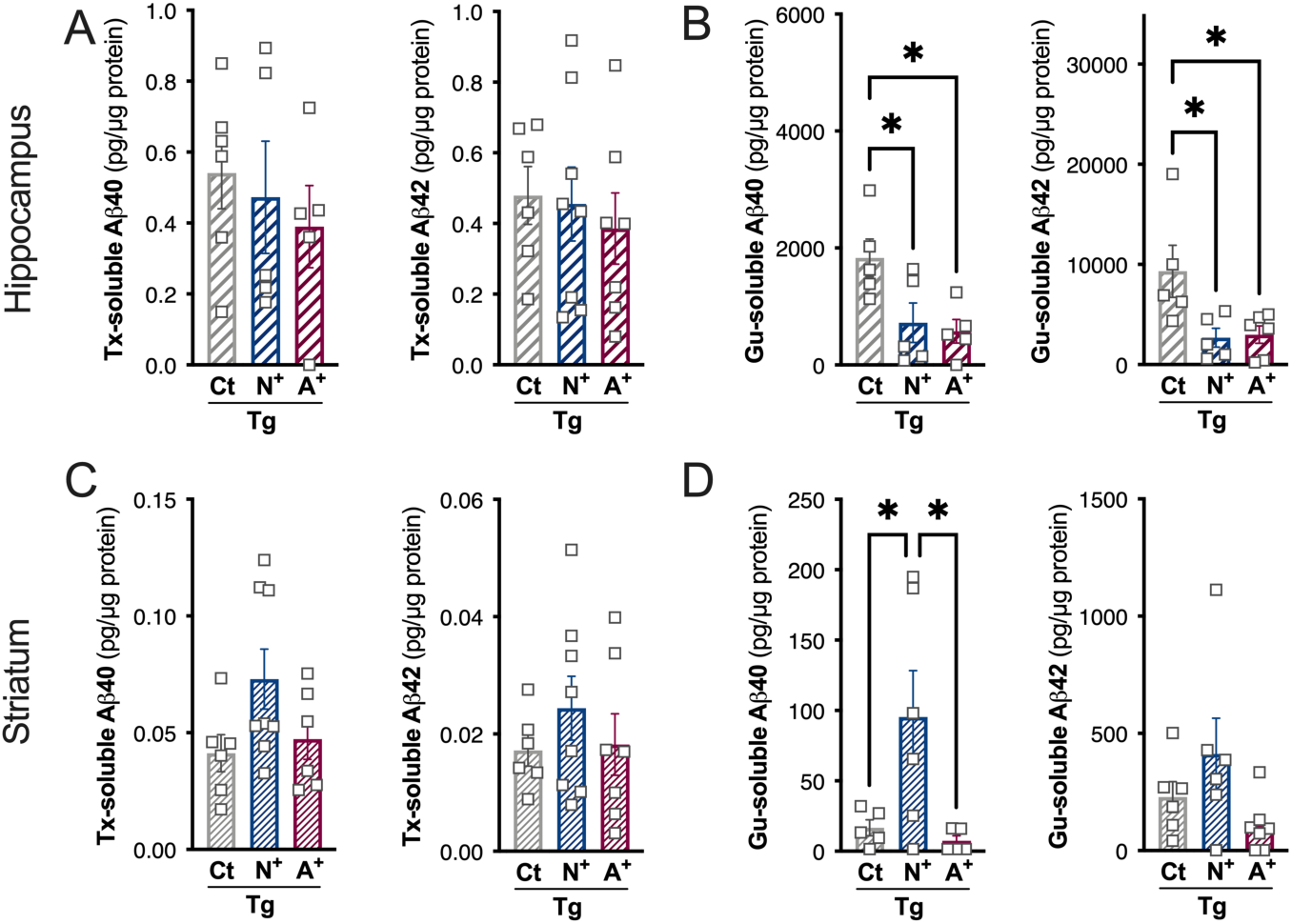
Chronic CNO stimulation of hippocampal astrocytes and neurons induces a reduction of the density of aggregated amyloid. **A.** Representative examples of AAV-N^+^ and AAV-A^+^ staining in the hippocampus. The AAV-N^+^ is present in neurons and absent from astrocytes (top panel) while the AAV-A^+^ is present in astrocytes and absent from neurons (lower panel). **B-C.** Quantitative analysis of Aβ40 and Aβ42 concentrations in triton- (Tx) and guanidine- (Gu) soluble fractions of proteins in the hippocampus. One-way ANOVA with the Fisher’s LSD posthoc test. *p<0.05 as compared to the Tg-Ct group. **D-E.** Quantitative analysis of Aβ40 and Aβ42 concentrations in triton- (Tx) and guanidine- (Gu) soluble fractions of proteins in the striatum. One-way ANOVA with the Fisher’s LSD posthoc test. *p<0.05 as compared to the Tg-Ct group.

In the striatum, Tx- and Gu-forms are also detected although their concentration is about 15 and 50 times lower than in the hippocampus, respectively. The only significant effect is the increase of Gu-Aβ40 in response to chronic stimulation of neurons (Fig. 6C-D, one-way ANOVA Gu-Aβ40, main effect of treatment: F2,13=5.05, p=0.023; one-way ANOVA Tx-Aβ40, main effect of treatment: F2,17=2.58, p>0.05; one-way ANOVA Tx-Aβ42, main effect of treatment: F2,18=0.66, p>0.05; one-way ANOVA Gu-Aβ42, main effect of treatment: F2,16=2.71, p>0.05).

### Chronic DREADD activation induced a lower glial cell reactivity in Tg rats

One index of glial reactivity is the measurement of the density of astrocytic and microglial cells. To determine the glial response to chronic stimulation of hippocampal neurons or astrocytes, an analysis of IBA1 and GFAP density was conducted. Fig 7A shows representative example of IBA1 and GFAP staining in the hippocampus. Chronic stimulation of neurons and astrocytes induced an overall increase in microglia density in the subiculum, dorsal and ventral hippocampus in WT animals (Fig. 7B, two-way ANOVA, main effect of AAV: F2,58=6.56, p=0.0027; main effect of area: F2,58=2.39, p=0.09; interaction: F4,58=0.9, p>0.05). In Tg rats, a trend or a significant increase in microglial density is observed in response to chronic stimulation of neurons and astrocytes, respectively (Fig. 7B, two-way ANOVA, main effect of AAV: F2,51=3.2, p=0.048; main effect of area: F2,51=1.64, p>0.05; interaction: F4,51=0.23, p>0.05). Chronic stimulation of neurons and astrocytes did not induce a difference in hippocampal astrocyte density in either WT or Tg (Fig. 7C, WT: two-way ANOVA, main effect of AAV: F2,58=1.72, p=0.18; main effect of area: F2,58=135.5, p<0.0001; interaction: F4,58=0.89, p>0.05).

**Figure 7.**
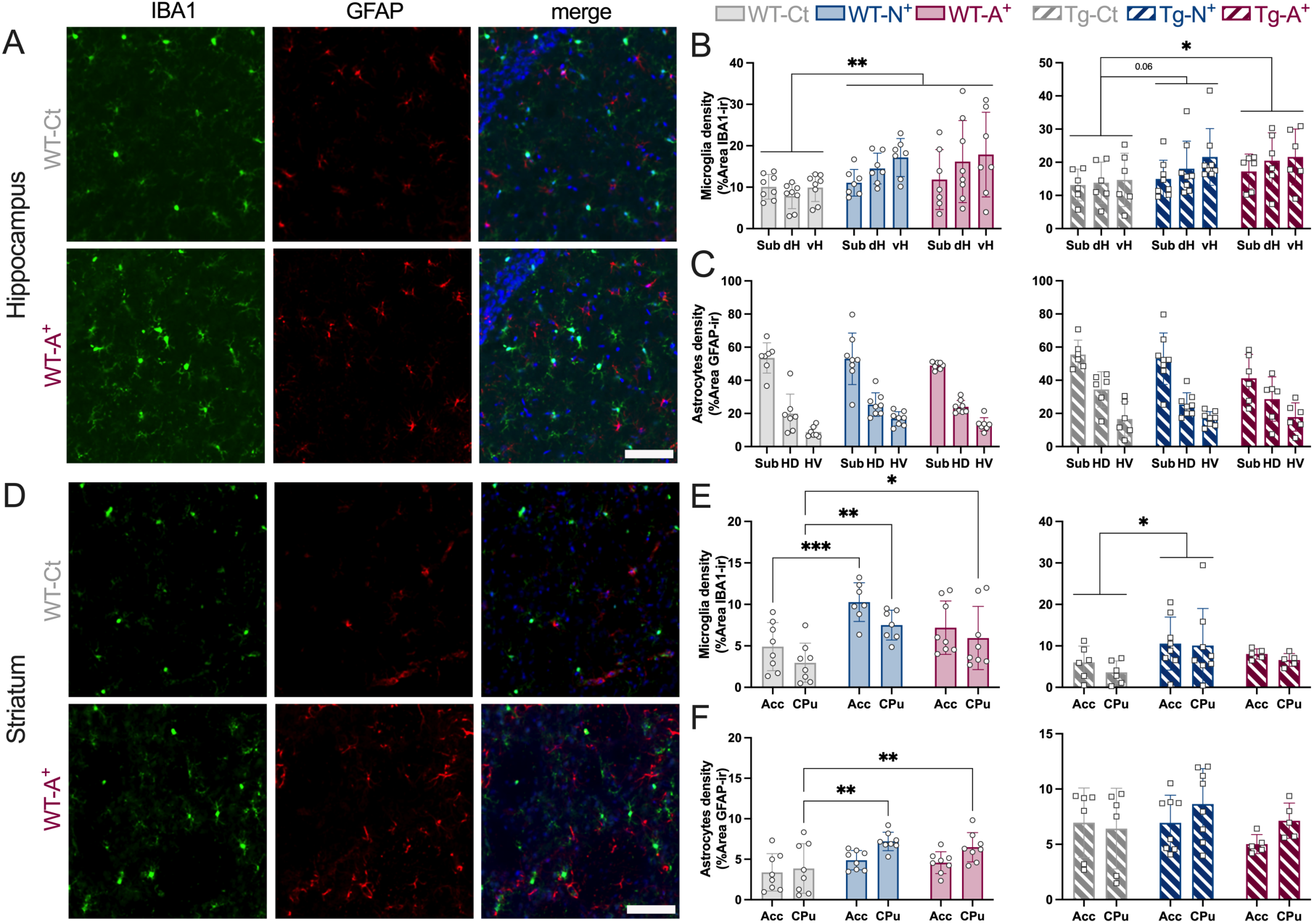
Glial cell activity in response to chronic CNO. **A.** Representative example of IBA1 and GFAP staining in WT-Ct and WT-A^+^ rats at the level of the dorsal hippocampus. **B.** Microglia density (% of IBA1^+^ area) in the hippocampal subdivisions (dH, dorsal hippocampus; sub, subiculum; vH, ventral hippocampus) in WT and Tg rats. Two-way ANOVA analysis with the Fisher’s LSD posthoc test. *p<0.05, **p<0.01, as compared to the respective Ct group. **C.** Astrocytes density (% of GFAP^+^ area) in the hippocampal subdivisions (dH, sub and vH) in WT and Tg rats. Two-way ANOVA analysis with the Fisher’s LSD posthoc test. **D.** Representative example of IBA1 and GFAP staining in WT-Ct and WT-A^+^ rats at the level of the caudate/putamen. **E.** Microglia density (% of IBA1^+^ area) in the striatum (accumbens, Acc; cadate/putamen, CPu) in WT and Tg rats. Two-way ANOVA analysis with the Fisher’s LSD posthoc test. *p<0.05, **p<0.01, ***p<0.001 as compared to the respective Ct group. **F.** Astrocytes density (% of GFAP^+^ area) in the hippocampal subdivisions (dH, sub and vH) in WT and Tg rats. Two-way ANOVA analysis with the Fisher’s LSD posthoc test. **p<0.01 as compared to the respective Ct group.

A representative example of IBA1 and GFAP staining in the striatum is given in Fig. 7D. The DREADD stimulation of hippocampal neuron induced an increase of microglial density in the Acc and the CPu, and the stimulation of hippocampal astrocytes elevated microglial density in the CPu (Fig. 7E, two-way ANOVA, main effect of AAV: F2,40=11.37, p=0.0001; main effect of area: F1,40=5.57, p=0.023; interaction: F2,40=0.27, p>0.05). In Tg rats, only neuronal stimulation induced an overall increase in microglia levels in the striatum (Fig. 7E, two-way ANOVA, main effect of AAV: F2,32=3.41, p=0.045; main effect of area: F1,32=0.61, p>0.05; interaction: F2,32=0.1, p>0.05).

Stimulation of hippocampal neurons and astrocytes induced an increase in astrocyte density in the CPu in WT but not in Tg rat (Fig. 7F, two-way ANOVA astrocytes in WT, main effect of AAV: F2,42=6.95, p=0.0025; main effect of area: F1,42=7.88, p=0.0075; interaction: F2,42=0.99, p>0.05; two-way ANOVA astrocytes in Tg, main effect of AAV: F2,32=1.29, p>0.05; main effect of area: F1,32=1.43, p>0.05; interaction: F2,32=0.78, p>0.05).

### Chronic DREADD activation stimulates phagocytosis of Aβ by glial cells

To determine whether the decrease in aggregated forms of amyloid is a consequence of stimulation of their degradation by glial cells, a measurement of the colocalization between dense MXO4^+^ plaques and glia markers (IBA1 and GFAP) was performed (see examples of staining in Fig. 8A). Stimulation of hippocampal neurons and astrocytes induced an increase in the density of amyloid plaques covered by microglial cells (Fig. 8B, two-way ANOVA, main effect of treatment: F2,411=4.55, p=0.011; main effect of distance from maximum coverage: F25,411=25.18, p<0.0001). Stimulation of hippocampal neurons but not astrocytes also induced an increase in the density of amyloid plaques covered by astrocytes (Fig. 8C, two-way ANOVA, main effect of treatment: F2,379=7.25, p=0.0008; main effect of distance from maximum coverage: F24,379=66.95, p<0.0001).

**Figure 8.**
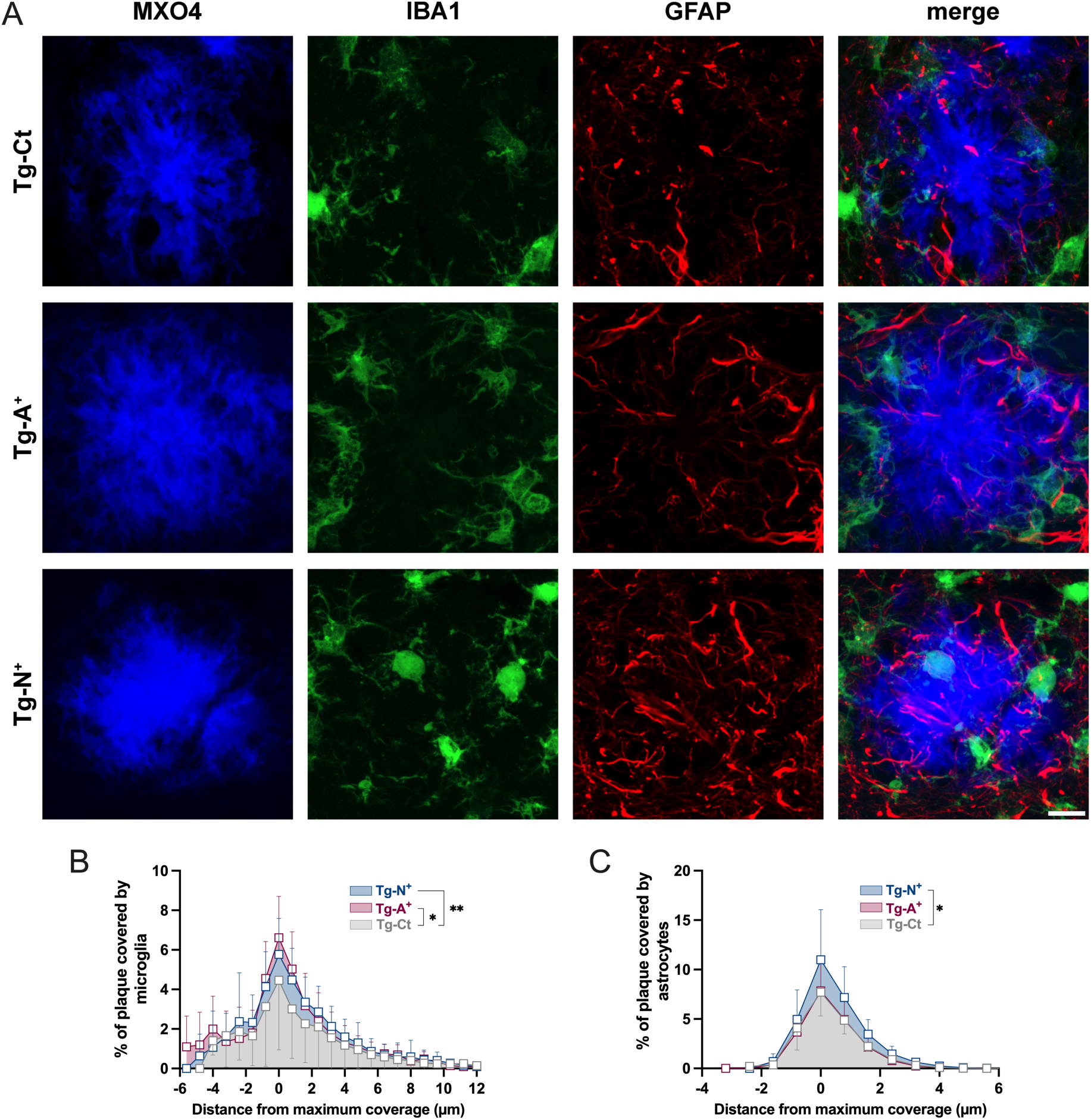
Chronic CNO stimulates amyloid phagocytosis. **A.** Representative example of IBA1^+^ and GFAP^+^ staining in MxO4^+^ area in the hippocampus of Tg rats. WT rats did not display any MxO4^+^ staining (not shown). **B.** % of Aβ plaque colocalized with microglia (IBA1^+^MxO4^+^ area in MxO4^+^ area) in the hippocampus. Two-way ANOVA analysis with the Dunnett’s multiple comparisons posthoc test. *p<0.05, **p<0.01, as compared to the Tg-Ct group. **C.** % of Aβ plaque colocalized with astrocytes (GFAP^+^MxO4^+^ area in MxO4^+^ area) in the hippocampus. Two-way ANOVA analysis with the Dunnett’s multiple comparisons posthoc test. *p<0.05 as compared to the Tg-Ct group.

## Discussion

In this study, we assessed the behavioural and neurochemical effects of acute and chronic neuronal and astrocytic stimulation using excitatory DREADD in a rat model of AD pathology. Our results show that acute and chronic neuronal and astrocytic stimulation induces widespread effects on the brain regional activation pattern, notably with an inhibition of striatal activation. This effect might be explained by the enhancement of dopamine release in the striatum by acute and chronic hippocampal neuronal and astrocytic stimulation. On the other hand, in a rat model of AD, these effects, especially of the chronic stimulation, are blunted, both in regional brain activation pattern and striatal dopaminergic signalling. The dysfunctional connectivity between the hippocampus and other brain regions is also supported by the finding that DREADD stimulation in the hippocampus increased microglial density and its capacity to limit AD pathology, whereas these effects were absent in the striatum.

Dysfunctional dopaminergic signalling in AD has been suggested by human molecular imaging studies^17–19^ and might be implicated in the occurrence of psychotic and other behavioural and psychological symptoms of the disease. Our work indicates that even in the absence of local AD pathology in the striatum (as was the case for our rats, which only had soluble Aβ forms and no lesions), hippocampal pathology may already induce circuit alterations that impact dopaminergic neurotransmission^20^. Chronic dopaminergic signalling in the striatum of WT rats induced a reduction in D_2_ receptor density, as expected. This effect is a compensatory property of neurotransmitter receptors upon chronic stimulation. The fact that in Tg rats the receptor density was unchanged support the notion that mesostriatal dopaminergic signalling does not respond to chronic hippocampal neuronal or astrocytic stimulation in the same way as WT rats. These observations support a presynaptic dopaminergic dysfunction, supported by the increased reactivity of postsynaptic receptors in the striatum^21^. This dopaminergic dysfunction may be contributing to the pathophysiology of psychotic syndromes in dementia, as is the case in other psychiatric pathologies involving psychotic symptoms^13^.

Chronic DREADD activation of astrocytes has been shown to induce transcriptomic changes in adjacent microglia, notably with the upregulation of a set of Alzheimer’s disease-associated genes that are associated to a microglial phenotype which is protective against AD pathology^31,32^. Our results extend this finding by showing that, functionally, the result of chronic astrocytic DREADD stimulation is associated to an increase in microglial density and especially in a microglial phenotype that more efficiently responds to and limits AD pathology. This effect was limited to the hippocampus, as chronic activation of neurons and astrocytes failed to reduce Aβ in the striatum, despite a significant increase in microglial density in this context and it even increased it. Indeed, chronic neuronal DREADD stimulation in the hippocampus induced an increase in the concentration of the Gu-soluble form of Aβ40 in the striatum. Moreover, a tendency towards an increase was observed for all forms of Aβ in this region. At the same time, the density of microglial cells was also increased. This discrepancy, i.e. the fact that microglial density increased in both the hippocampus and the striatum after chronic DREADD stimulation, but Aβ is reduced in the hippocampus and increased in the striatum could be explained by the effect of dopamine directly on microglia. Indeed, previous studies have shown that dopamine, through various receptors, can reduce proinflammatory activation and promote an overall protective phenotype. We may thus hypothesize that the blunting of the dopaminergic signalling to the striatum in Tg rats, deprives microglial cells of this protective effect of dopamine and fails to respond to AD pathology efficiently^33,34^. These results suggest that dysfunction of the dopaminergic circuitry in AD could also impact microglial phenotype. This could be particularly relevant for the understanding of the pathophysiology of syndromes such as AD psychosis, in which dopaminergic dysfunction is implicated^35^.

The fact that neuron and astrocyte stimulation locally induced a microglial phenotype that showed improved performance against Aβ pathology may also have important implications for strategies involving brain stimulation to reduce Aβ load and limit the disease progression^36–41^. Our results support these approaches, although they indicate that the altered connectivity of the hippocampus with other brain regions may spatially limit the effect of neuronal and astrocytic stimulation to the areas that are being directly stimulated. Nevertheless, if brain stimulation strategies are applied to a sufficiently early time-point of the AD pathological process, when Aβ is present but has not yet induced dysfunctional connectivity, a locally applied stimulation in the hippocampus can potentially have brain-wide protective effects. Future work is needed to better characterize AD connectivity changes in the trajectory of AD pathology and inform on the optimal timing of the application of such approaches.

We recognize that our study has limitations. While we make a number of novel observations, a more thorough assessment of the phenotypic alterations in neurons and glial cells after acute and chronic DREADD stimulation is needed. In addition, although the potential to alter mesostriatal transmission via optogenetic or chemogenetic stimulation of the hippocampus is known^23^, further study should clearly delineate the exact circuitry that is implicated in this phenomenon and shows vulnerability towards AD pathology. Finally, we only used male rats and these findings need to be reproduced in female animals, given the impact of sex on AD pathology itself and on the phenotype of glial cells^42^.

In conclusion, our work suggests that hippocampal neuronal and astrocytic stimulation have widespread effects on regional rat brain activity. Hippocampal neuronal and astrocytic activation also stimulates dopaminergic signalling to the striatum both acutely and chronically, although these brain-wide and striatum-specific effects are blunted in a rat model of AD pathology. Stimulation of neurons and astrocytes in the hippocampus locally stimulates a protective, Aβ-limiting phenotype of microglia, which surrounds Aβ plaques and limits Αβ concentration more efficiently.

## Acknowledgments

This work was supported by the Swiss National Science Foundation (grants 186517 and 184713).

