## Supplemental table 1 for "Impairment of hippocampal astrocyte-mediated striatal dopamine release and locomotion in Alzheimer’s disease"

|  | **WT** |  |  | **TgF344-AD** |  |  |
| --- | --- | --- | --- | --- | --- | --- |
| Amygdala | 81.94 | **±** | 1.93 | 84.73 | **±** | 4.30 |
| Bed Nucleus Stria Terminalis | 101.01 | **±** | 1.98 | 100.82 | **±** | 2.61 |
| Caudate Putamen | 104.80 | **±** | 4.19 | 104.23 | **±** | 6.42 |
| Cerebellum | 121.63 | **±** | 4.14 | 116.83 | **±** | 6.97 |
| Cortex Auditory | 89.81 | **±** | 1.48 | 89.37 | **±** | 3.79 |
| Cortex Cingulate | 106.88 | **±** | 7.38 | 111.50 | **±** | 2.66 |
| Cortex Piriform | 81.66 | **±** | 1.25 | 87.47 | **±** | 3.90 |
| Cortex Frontal Association | 88.68 | **±** | 4.05 | 91.47 | **±** | 8.54 |
| Cortex Insular | 89.30 | **±** | 1.99 | 93.27 | **±** | 5.80 |
| Cortex Medial Prefrontal | 111.88 | **±** | 3.52 | 116.83 | **±** | 2.48 |
| Cortex Motor | 93.19 | **±** | 2.77 | 94.70 | **±** | 3.74 |
| Cortex Orbitofrontal | 103.23 | **±** | 4.33 | 107.30 | **±** | 11.23 |
| Cortex Parietal Association | 88.90 | **±** | 2.23 | 89.08 | **±** | 0.95 |
| Cortex Retrosplenial | 112.50 | **±** | 3.25 | 115.17 | **±** | 7.57 |
| Cortex Somatosensory | 90.39 | **±** | 2.11 | 89.75 | **±** | 4.12 |
| Cortex Temporal Association | 90.14 | **±** | 2.15 | 93.60 | **±** | 2.58 |
| Cortex Visual | 99.75 | **±** | 4.27 | 104.17 | **±** | 3.06 |
| Globus Pallidus | 98.71 | **±** | 4.93 | 96.50 | **±** | 5.06 |
| Medial Geniculate | 105.50 | **±** | 4.14 | 104.12 | **±** | 6.54 |
| Medulla | 91.75 | **±** | 3.04 | 88.78 | **±** | 4.08 |
| Mesencephalic Region | 105.25 | **±** | 3.49 | 103.22 | **±** | 4.11 |
| Olfactory Nuclei | 90.56 | **±** | 6.49 | 92.77 | **±** | 15.09 |
| Periaqueductal gray | 118.75 | **±** | 5.06 | 114.83 | **±** | 5.74 |
| Pons | 83.19 | **±** | 2.39 | 81.82 | **±** | 3.79 |
| Septum | 108.50 | **±** | 5.42 | 106.33 | **±** | 5.24 |
| Superior Colliculus | 125.75 | **±** | 3.54 | 119.83 | **±** | 3.06 |
| Accumbens | 106.38 | **±** | 2.26 | 108.83 | **±** | 5.38 |
| Hippocampus | 108.75 | **±** | 1.04 | 108.50 | **±** | 3.56 |
| Hypothalamus | 93.06 | **±** | 1.60 | 95.52 | **±** | 8.72 |
| Thalamus | 104.88 | **±** | 1.73 | 101.23 | **±** | 3.54 |

**Supplemental Table 1. The [^99^Tc]HMPAO brain cartography is similar between WT and TgF344-AD rats.**

Values are expressed as mean of the left/right region and normalized to the whole brain ± SD**.**
